# Multi-Site Investigation of Gut Microbiota in CDKL5 Deficiency Disorder Mouse Models: Targeting Dysbiosis to Improve Neurological Outcomes

**DOI:** 10.1101/2024.03.19.581742

**Authors:** Francesca Damiani, Maria Grazia Giuliano, Sara Cornuti, Elena Putignano, Andrea Tognozzi, Vanessa Suckow, Vera M. Kalscheuer, Paola Tognini

## Abstract

**Background:** Cyclin-Dependent Kinase-Like 5 (CDKL5) deficiency disorder (CDD) is a rare X-linked developmental encephalopathy caused by pathogenic variants of the CDKL5 gene. In addition to a diverse range of neurological symptoms, CDD patients frequently manifest gastrointestinal (GI) issues and subclinical immune dysregulation. This comorbidity suggests a potential association with the intestinal microbiota, prompting an investigation into whether gut dysbiosis contributes to the severity of both GI and neurological symptoms.

**Methods:** We examined the gut microbiota composition in two CDKL5 null (KO) mouse models in males at three different developmental stages: postnatal day (P) 25 and P32 during youth, and P70 during adulthood.

**Results:** Changes in diversity and composition were observed, particularly during juvenile ages, suggesting a potential gut microbiota dysbiosis in the CDD mouse models. To further understand the role of the gut microbiota in CDD, we administered an antibiotic cocktail to the mice and conducted functional and behavioral assessments. Remarkably, significant improvement in visual cortical responses and reductions in hyperactive behavior were observed. To shed light on the cellular mechanisms we focused on microglia. Alterations in specific aspects of microglia morphology, indicative of activation state and surveillance of the microenvironment, were observed in the CDKL5 KO mice and ameliorated by antibiotic administration.

**Conclusions:** Our findings highlight the potential impact of modifications in the intestinal microbiota on the severity of CDD symptoms, expanding our understanding beyond GI disturbances to encompass influences on neurological outcomes. This cross-border study provides valuable insights into the intricate interplay between gut microbiota and neurodevelopmental disorders.

## INTRODUCTION

Cyclin-Dependent Kinase-Like 5 (CDKL5) deficiency disorder (CDD) is an X-linked early-onset encephalopathy caused by mutations in the CDKL5 gene. The estimated prevalence of this condition is 1 in 40,000 to 1 in 60,000 live births. Patients with CDD exhibit severe global developmental delays, including early-onset epileptic encephalopathy, intellectual disability, visual and motor impairments, strong hypotonia, sleep disturbances, and hand stereotypies (1–3). Some of these symptoms overlap with Rett Syndrome (RTT); however, the key distinctions include the absence of a regression period and the early onset of seizures in CDD patients. Currently, there are no available treatments for CDD.

A common issue in patients with CDD is the presence of gastrointestinal (GI) problems, including diarrhea, constipation, and acidic reflux (4). This may be linked to subclinical immune dysregulation, likely stemming from a faulty inflammation regulatory signaling system (5,6). Remarkably, GI disorders are about four times more common in children with autism spectrum disorders (ASD) than in neurotypical children (7). Also, other central nervous system (CNS)-related comorbidities, such as seizures and sleep disorders, tend to occur more frequently in individuals with GI dysfunction in the ASD population (8–10). The heightened occurrence of GI dysfunction in ASD, coupled with its substantial correlation with challenging behaviors and psychiatric comorbidities, posits a plausible association between gut and brain dysfunction in CDD. At least part of this association could be due to abnormal composition of the gut microbiome, i.e. the assemblage of microorganisms residing in the GI tract that engage symbiotic interactions with their host (11,12). Indeed, ASD patients display alterations in the composition of the gut microbiota (13) as well as RTT patients (14,15). Importantly, we have recently discovered differences in biodiversity and composition of intestinal microbes in a cohort of Italian patients affected by CDD (16). Disturbances in the composition or equilibrium of the gut microbiota, termed dysbiosis, have been associated with inflammatory bowel disease (IBD), metabolic disturbances, and even neuropsychiatric conditions (17–20). Nevertheless, the composition of the intestinal microbiota in widely used CDD models, and the impact of alterations in microbiota composition on neurologically relevant outcomes has not been explored.

To investigate the link between CDD symptoms and the gut-brain axis, we analyzed the developmental trajectory of the fecal microbiota in CDKL5 mouse models at distinct ages, and found significant differences compared to wild type (WT) littermates. Since the intestinal microbiota is strongly affected by environmental factors, our investigation was performed considering two CDKL5 null mouse models, living in different facilities in Italy and Germany. Our findings unveiled the presence of dysbiosis, suggesting a potential negative impact of the gut microbiota of CDKL5 mutants on neurological outcomes. To test this hypothesis we targeted the gut microbiota by a 25-day treatment with an antibiotic cocktail (ABX) and we assessed visual cortical responses and behavioral performance. Moreover, we analyzed microglia alterations as a possible cellular underpinning of the ABX effect. The results revealed that ABX intervention partially rescued this phenotype, indicating an improvement in the potential inflammatory state of the brain in CDD. Overall, our study shows that microbial manipulation could be a novel and non-invasive strategy to improve symptoms in CDD patients.

## MATERIALS AND METHODS

### Animals

All experiments were carried out in accordance with the European Directives (2010/63/EU), and were approved by the Italian Ministry of Health.

Animals were kept in rooms at 22°C with a standard 12 h light–dark cycle. During the light phase, a constant illumination below 40 lux from fluorescent lamps was maintained. Food (standard diet, 4RF25 GLP Certificate, Mucedola) and water were available ad libitum and changed weekly. The mice from the National Research Council vivarium in Pisa and used in this work derive from the Cdkl5 null strain in C57BL/6N background developed in (21), and backcrossed into C57BL/6J over seven generations. The mice from the Berlin vivarium used to characterize the fecal microbiota composition were derived from a conditional mutant mouse line crossed to C57BL/6J *Hprt* Cre-deleter to generate the Cdkl5 KO mice with exon 4 deletion (manuscript in preparation).

Male wild type (WT) mice were bred with heterozygous female mice to obtain mutant and WT littermates. Weaning was performed on postnatal day (P)21–23, and mice were individually housed.

In the current work we used male CDKL5-/y (CDKL5 KO) mice and male CDKL5+/y (WT) littermates.

Data analyses were performed by experimenters blind to the animal genotype.

### Fecal DNA extraction and 16S rRNA sequencing

To analyze the composition of the fecal microbiota of CDKL5 KO and WT littermate mice at different ages, fresh feces were collected longitudinally from the same subjects at P25, P32 and P70, snap-frozen in liquid nitrogen and stored at -80 °C. To avoid the cage-effect and litter effects on microbiota composition, the animals used for the analysis belonged to different cages and different litters both in Pisa and Berlin vivariums. Bacterial DNA was extracted using the QIAamp Powerfecal DNA kit (Qiagen), and its concentration was quantified by Nanodrop 2000 C Spectrophotometer (ThermoFisher Scientific).

The 16S rRNA sequencing and analysis was performed by a service offered by Institute of Applied Genomics (IGA, Udine, Italy). For details see “Supplementary Methods”.

### Antibiotic treatment

CDKL5 KO mice were treated with an antibiotic cocktail (ABX, vancomycin 0.5 g/l, ampicillin 1 g/l and neomycin 1 g/l) dissolved in drinking water from weaning (P21) to P45. The ABX was refreshed every other day. Control WT littermates and control CDKL5 KO mice were weaned at P21 and drank regular water until P45. At the end of treatment mice were transcardially perfused.

### Behavioral analysis

#### Hindlimb clasping

Mice were suspended by their tail for 2 min and hindlimb clasping was assessed from video recordings. A lower clasping score was assigned if the mouse maintained the corresponding hindlimb clasping position for a minimum of 8 seconds during the observation period. The scoring system was adapted from (22). Specifically, a score of 4 was assigned if both hindlimbs consistently remained fully spread from the abdomen. If one hindlimb was retracted or both hindlimbs were partially retracted toward the abdomen without touching it, a score of 3 was assigned. If both hindlimbs were retracted toward the abdomen without touching each other, a score of 2 was given. If both hindlimbs were retracted, touching the abdomen and each other in a full clasping position, a score of 1 was assigned.

#### Nesting behavior

Mice were placed in a clean cage with two untouched nestlet (7 cm diameter rounds of pressed cotton) in the morning. The quality of the nest was scored 24 hours later by mouse genotype-blind observers. The nests were evaluated on a 5-point scale as described in (23).

#### Exploratory and Hyperactive Behavior, Y maze

A Y-shaped maze with three symmetrical gray solid plastic arms at a 120-degree angle (26 cm length, 10 cm width, and 15 cm height) was used to test exploratory and hyperactive behaviors. Mice were placed in the center of the maze and allowed to freely explore the maze for 8 minutes. The apparatus was carefully cleaned with ethanol between trials to avoid the build-up of odor traces. All sessions were video-recorded (Noldus Ethovision XT) for offline blind analysis. The arm entry is defined as all four limbs within the arm. A triad is defined as a set of three arm entries, when each entry is to a different arm of the maze. The number of arm entries and the number of triads were recorded in order to calculate the alternation percentage (generated by dividing the number of triads by the number of possible alternations and then multiplying by 100).

### Statistical Analysis

The sample sizes were based on prior studies and are indicated in the figure legend for each panel. Whenever possible, quantification and analyses were performed blind to the experimental condition. The majority of statistical analyses were performed using GraphPad Prism version 7 (GraphPad Software, San Diego, CA, USA). All data are represented as the mean ± SEM unless otherwise stated. N’s represent single animals unless otherwise stated. Statistical significance was defined in the figure panels as follows: *P≤0.05, **≤0.01 and **P≤0.001.

One-Way ANOVA or Kruskal-Wallis tests were applied when more than 2 experimental groups were analyzed. PERMANOVA was used to calculate statistics of beta diversity. See “Supplements” for details.

## RESULTS

### Alterations in the composition of CDKL5 KO fecal microbiota

To investigate the composition of intestinal bacteria, fresh fecal samples were collected longitudinally from both CDKL5 KO and WT mice. Sampling occurred longitudinally at postnatal day 25 (P25), again at P32, and during adulthood at P70 (see Fig. 1a). Bacterial DNA was then extracted from these fecal samples and subjected to analysis using 16S rRNA sequencing (seq).

**FIG.1.**
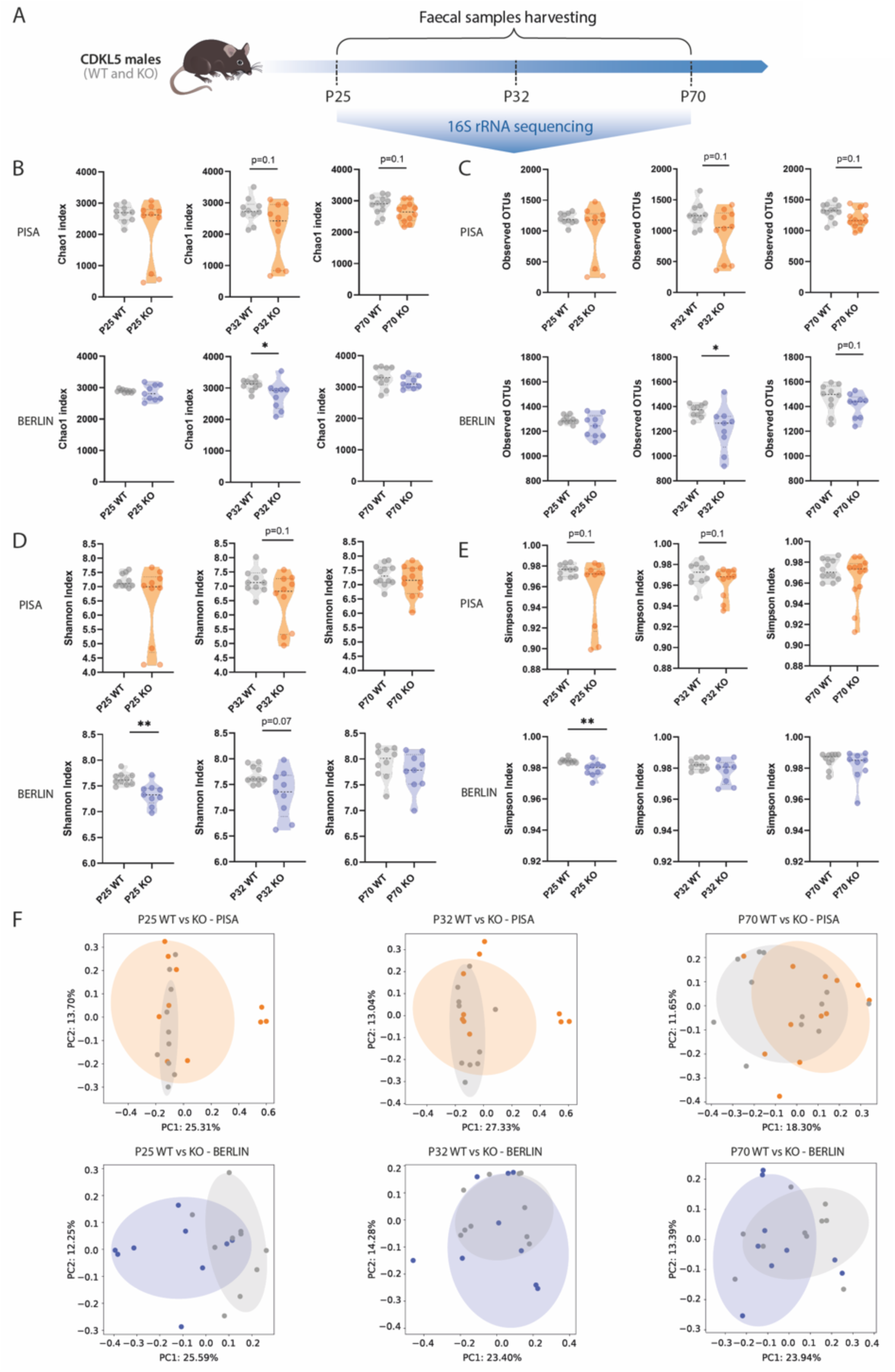
Longitudinal characterization of the gut microbiota in CDKL5 null mice raised in Pisa or Berlin vivaria. **(A)** Timeline for fecal samples harvesting. Mice were single-housed starting from weaning onwards and fecal samples were collected at postnatal day (P)25, P32 and P70 for gut microbiota characterization. **(B-E)** Violin plots showing alpha diversity comparison between WT and KO mice from Pisa (top plots) and Berlin (bottom plots) vivaria: **(A)** CHAO1 index, **(B)** Observed OTUs, **(C)** Shannon index, **(E)** Simpson index. Each dot represents a single animal. Median alpha diversity is shown as black dotted horizontal line. (n=9-12 mice per group, Mann-Whitney U-test, *p ≤ 0.05, **p < 0.01). **(F)** PCoA plot based on Bray-Curtis dissimilarity matrix showing beta diversity of WT and KO mice from Pisa (top plots) and Berlin (bottom plots) vivaria. The ellipses represent 95% confidence intervals for each group. Axes in the PCoA display the percentage of variation explained using Bray-Curtis dissimilarity (PERMANOVA test; mice from PISA: P25 KO versus P25 WT pseudo-F=1.718 p=0.058, P32 KO versus P32 WT pseudo-F=1.839 p=0.038, P70 KO versus P70 WT pseudo-F=1.421 p=0.097; mice from BERLIN: P25 KO versus P25 WT pseudo-F=1.817 p=0.002, P32 KO versus P32 WT pseudo-F=1.060 p=0.345, P70 KO versus P70 WT pseudo-F=1.053 p= 0.332).

Given that the composition of gut microbiota can be influenced by environmental factors, including the animal’s living conditions (i.e., the vivarium), we conducted our analysis on two different CDKL5 null mouse lines born and raised in two distinct animal facilities: the vivarium at the National Research Council in Pisa, Italy, and the vivarium at the Max Planck Institute for Molecular Genetics in Berlin, Germany. This parallel analysis of CDD mouse models from different facilities served to increase the robustness of the differences observed in our experiments.

Bacterial relative abundance is reported in Suppl. Table 1 (Pisa Vivaium) and Suppl. Table 2 (Berlin Vivarium).

To analyze the results, we first calculated alpha diversity, a parameter reflecting species richness or evenness in a microbial ecosystem (24). A high alpha diversity is generally associated with a healthy microbiome condition, while a reduction has been observed in a variety of diseases such as obesity (25), colitis, IBD (26), and to some extent neurological disorders (27). We calculated four different indexes and found a significant decrease in the Chao1 index (Fig. 1b) and observed OTU (Fig. 1c) at P32 in the CDKL5 KO raised in the Berlin vivarium. This result was paralleled by a tendency toward a significant decrease in alpha diversity in CDKL5 KO mice from Pisa vivarium (Fig.1b-c).

Moreover, Shannon Index and Simpson Index were decreased in the CDKL5 KO mice at P25 with respect to WT littermates in the Berlin Vivarium (Fig.1d-e). Second, we computed beta diversity, which measures the degree of phylogenetic similarity between microbial communities, using the Bray-Curtis dissimilarity (28). The principal coordinates analysis (PCoA) of the distance matrix of the fecal microbiota belonging to all the animals raised in the Pisa vivarium with respect to the ones raised in Berlin showed, as expected, a distinct clustering of the samples (Suppl. Fig.1). PCoA at the 3 different ages demonstrated the presence of a certain degree of clustering of fecal microbiota in WT animals versus CDKL5 KO mice. Specifically, significant phylogenetic dissimilarities were present between CDKL5 KO mice and WT littermates at P32 in the Pisa vivarium and at P25 in the Berlin vivarium (Fig. 1f).

Third, we sought to explore differences in the bacterial composition between juvenile and adult WT and CDKL5 KO mice by using Linear discriminant analysis (LDA) effect size (LEfSe) method (29). Lefse analysis revealed taxonomic dissimilarities between WT and CDKL5 KO mice in both the Pisa and Berlin facilities (Fig. 2a,c, e and Fig. 2b, e). In the Pisa vivarium, such differences were more pronounced during juvenile ages (P25 and P32 Fig. 2a,b) with respect to P70 (Fig. 2c, See also Suppl. Fig. 2). In particular, the family *Alcaligenaceae*, and the orders Burkholderiales and Pasteurellales were enriched in the CDKL5 KO microbiota at both P25 and P32. Together with *Enterobacteriaceae* and

**FIG.2.**
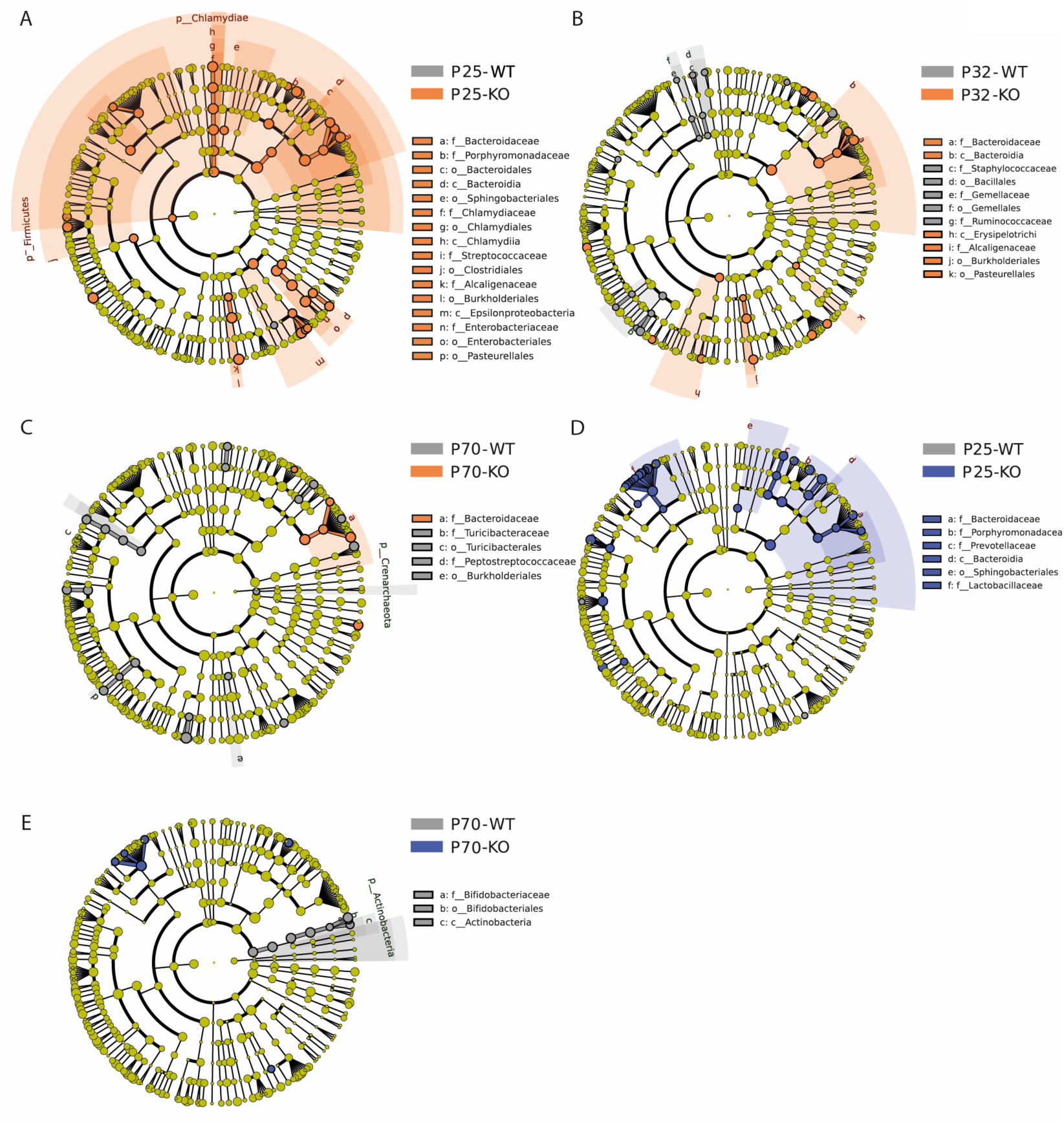
Linear discriminant analysis of Effect size of the fecal microbiota in WT vs CDKL5 KO mice. Taxonomic cladograms resulted from LEfSe analysis of the gut microbiota, showing significantly differentially enriched taxa (relative abundance ≥ 0.5%) for taxonomy level L7 (species) between WT and KO mice, at different ages, in Pisa and Berlin facilities. The colors represent the genotype in which the indicated taxa is more abundant with respect to the other genotype. Differential taxa were determined based on an LDA threshold score of >2.5 **(A-C)** Taxonomic differences between WT and CDKL5 KO mice from the Pisa vivarium, at P25 **(A)**, P32 **(B)** and P70 **(C)**. **(D-E)** Taxonomic differences between WT and KO mice from the Berlin vivarium, at P25 **(D)** and P70 **(E)**. No differences were observed at P32 for the mice living in the Berlin facility.

Epsilonproteobacteria enriched only in the P25 CDKL5 KO microbiota (Fig. 2a), those taxa belong to the phylum Proteobacteria. Emerging evidence identified the Proteobacteria (or Pseudomonadota) phylum as a possible microbial signature of disease (30,31). Notably, the family *Bacteroidaceae* were significantly enriched in the fecal microbiota of CDKL5 KO mice with respect to WT littermates at all ages (Fig. 2a-c). In the Berlin facility, the cladogram displayed a significant enrichment of the family *Porphyromodanaceae*, *Prevotellaceae*, *Lactobacillaceae*, in the class Bacteroidia, in the order Sphingobacteriales and again in the family *Bacteroidaceae* in the P25 CDKL5 KO mouse vs WT littermates (Fig. 2d), suggesting that the family *Bacteroidaceae* is a potential signature of the CDD mouse gut microbiota. While at P32 the LEfSe showed no differences, at P70 WT mice showed an enrichment of taxa related to intestinal health and probiotics, such as *Bifidobacteriaceae*, with respect to CDKL5 KO mice (Fig. 2e). Notably, *Bifidobacterium longum*, considered a probiotic species, was significantly decreased at P70 in both the Pisa and Berlin CDKL5 KO mice (Fig. 2c,e).

To further explore the taxa alteration signature of CDKL5 KO vs WT littermates gut microbiota, we subtracted the variable “vivarium” in the LEfSe analysis, grouping together WT and CDKL5 KO mice from Pisa and Berlin. At P25, CDKL5 KO mice displayed a significant enrichment in TAXA belonging to the order Burkholderiales, family *Alcaligenaceae*, and in *Bacteroidales*. The genera *Sutterella*, *Lactobacillus* and *Ruminococcus* were also taxa characterizing the CDKL5 KO microbiota (Fig. 3a and Suppl. Fig. 3). At both P25 and P32 *Bacteroides rodentium*, belonging to the family *Bacteroidaceae*, was enriched in the CDKL5 KO microbiota (Fig. 3b). Finally, the microbiota of CDD mice seem to be defective in the genera *Allobaculum* and *Parapedobacter* at P70 compared to WT littermates (Fig. 3c).

**FIG.3.**
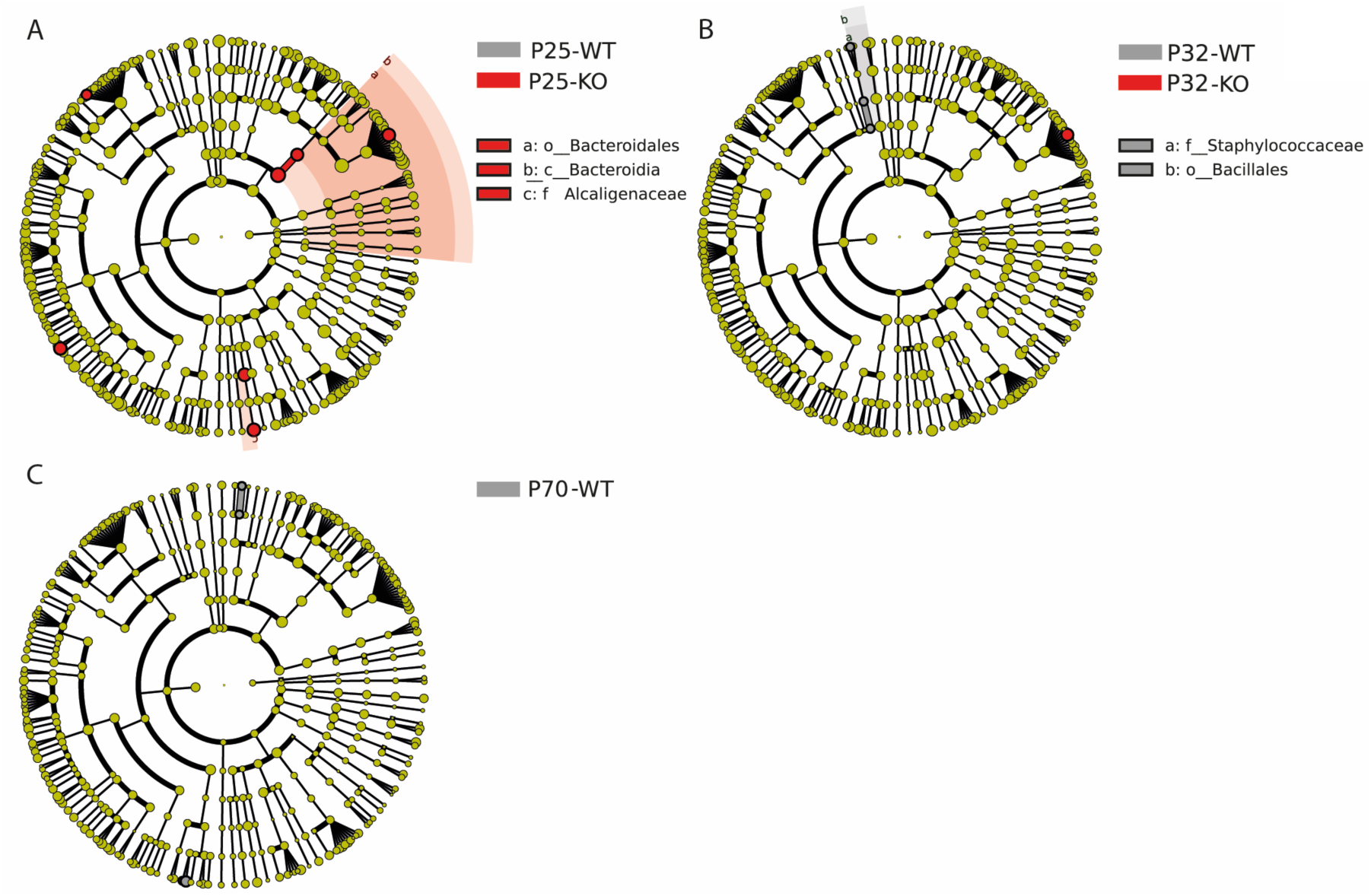
Linear discriminant analysis of Effect size of the fecal microbiota in WT vs CDKL5 KO mice subtracting the variable mouse facility. Taxonomic cladograms resulted from LEfSe analysis of the gut microbiota, showing significantly differentially enriched taxa (relative abundance ≥ 0.5%) for taxonomy level (species) between WT and KO mice, at different ages, subtracting the variable mouse facility.

Although, as expected, the differences between CDKL5 KO mice and WT littermates in the Pisa and Berlin vivarium did not perfectly overlap, our analysis highlighted significant changes in specific signature taxa in both mouse facilities, pointing toward stronger differences at younger ages.

Overall, our findings provide valuable insights into the dynamic relationship between gut microbiota composition and CDKL5 deficiency, shedding light on potential microbial signatures associated with this disorder across different developmental stages and environmental settings.

### Antibiotic treatment improves functional and behavioral outcomes in CDKL5 KO mice

As the composition analysis revealed differences in the CDKL5 fecal microbiota, we hypothesize the presence of a dysbiosis condition. Dysbiosis has been observed in several neuropsychiatric disorders (32), and in mouse models has been linked to behavioral and neurofunctional impairments (11,33). Therefore, we treated mice with an antibiotic cocktail (ABX) in drinking water to target intestinal microbes, particularly bacteria taxa, and tested the potential improvement in functional and behavioral outcomes. As prominent differences in the fecal microbiota were observed in the CDKL5 null mouse at younger ages in both the Pisa and Berlin facilities, mice were subjected to ABX administration from weaning (P21) for the next 24 days (Fig. 4a).

**FIG.4.**
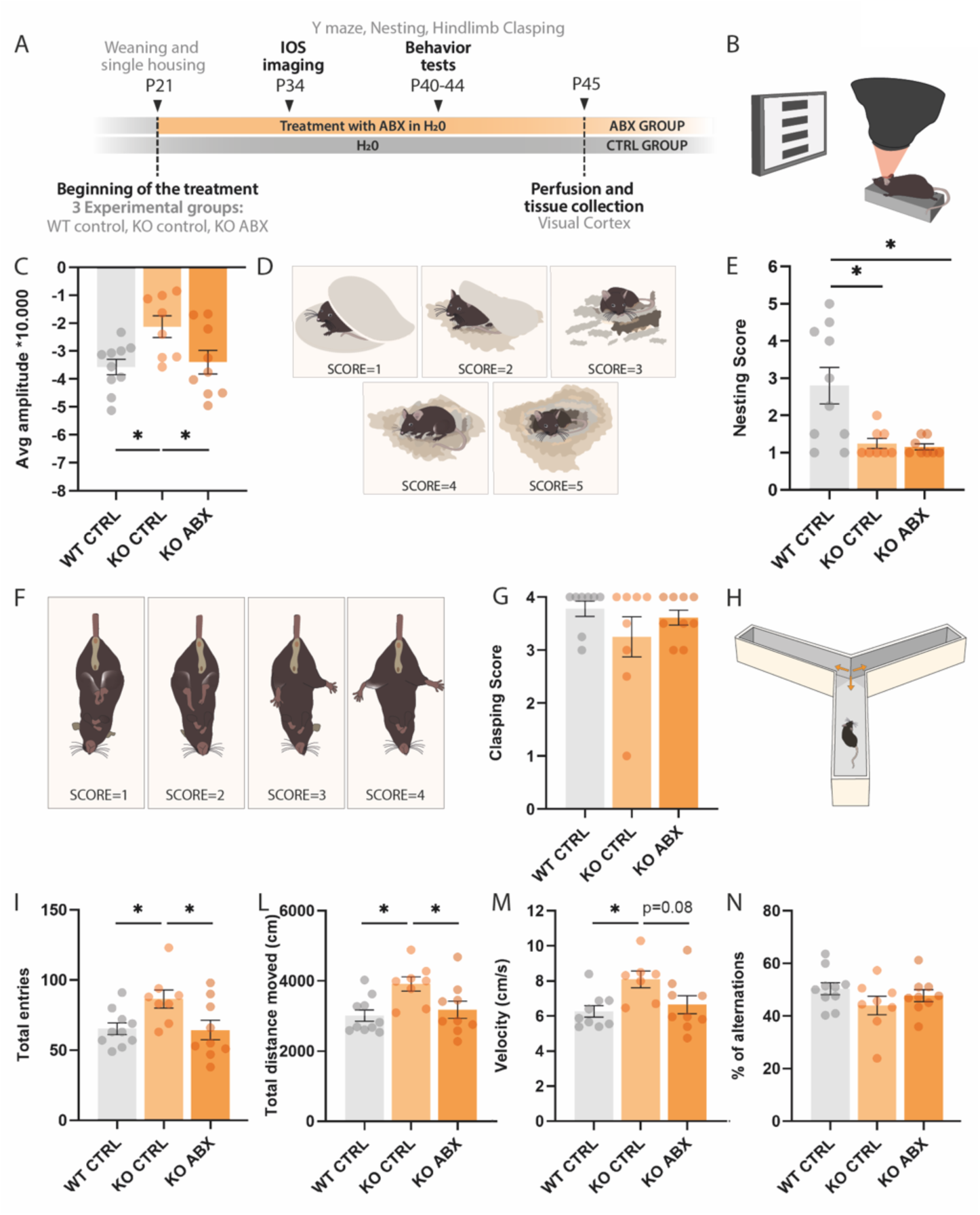
ABX administration significantly improves visual cortical responses and reduces the hyperactive behaviour of KO mice. **(A)** Experimental timeline. Mice were single housed starting from weaning (P21) onwards. A battery of behavioural and functional tests were performed in three experimental groups (n=8-10 mice per group): WT control group (WT CTRL), CDKL5 KO control group (KO CTRL), CDKL5 KO ABX-treated group (KO ABX). **(B)** Schematic illustration of the setup for the IOS imaging. **(C)** The graph represents the average amplitude of the cortical responses to contralateral eye stimulation (One-way ANOVA p=0.021, multiple comparisons Tukey post-hoc test; WT CTRL versus KO CTRL p=0.025, WT CTRL versus KO ABX p=0.936, KO CTRL versus KO ABX p=0.058). **(D)** Schematic representation of the scoring system for nest building assessment. **(E)** Nest building ability scored at 24 hour after placement of the untouched nestlet (Kruskal-Wallis test p=0.014, Dunn’s post-hoc test; WT CTRL versus KO CTRL p=0.057, WT CTRL versus KO ABX p=0.028, KO CTRL versus KO ABX p>0.999). **(F)** Schematic representation of the scoring system to evaluate hindlimb clasping behavior. **(G)** Scoring obtained upon 2 minutes of tail suspension (Kruskal-Wallis test p=0.467). **(H)** Schematic representation of the arena used for the Y-maze task. **(I)** Y maze number of total entries (One-way ANOVA p=0.024, multiple comparisons Tukey post-hoc test; WT CTRL versus KO CTRL p=0.043, WT CTRL versus KO ABX p=0.992, KO CTRL versus KO ABX p=0.038). **(L)** Y maze total distance moved in centimetres (cm) (One-way ANOVA p=0.012, multiple comparisons Tukey post-hoc test; WT CTRL versus KO CTRL p=0.013, WT CTRL versus KO ABX p=0.830, KO CTRL versus KO ABX p=0.052). **(M)** Velocity (cm/s) in the Y-maze (One-way ANOVA p=0.026, multiple comparisons Tukey post-hoc test; WT CTRL versus KO CTRL p=0.025, WT CTRL versus KO ABX p=0.808, KO CTRL versus KO ABX p=0.085). **(N)** % of alterations in the arms (One-way ANOVA p=0.725). Error bars represent SEM. Circles represent single experimental subjects.

As the gut bacteria are involved in energy harvesting from the ingested food, we monitored the body weight of the treated mice. We found a slight, albeit significant reduction in the body weight (Suppl. Fig. 4a). In addition, we investigated the efficacy of ABX treatment, and found a significant decrease of bacterial DNA concentration in the feces of CDKL5 KO mice after ABX administration compared to controls (Suppl. Fig. 4b). Both CDD patients and CDKL5 null mice present a visual deficit (2,34–37). After 13 days of ABX administration, we assessed the visual cortical responses by optical imaging of the intrinsic signal (IOS) (Fig. 4b). As expected, CDKL5 KO controls (CTRL) that drank normal water showed a significant reduction in the amplitude of visual responses compared to WT littermate controls (Fig. 4c). Remarkably, ABX administration rescued the visual response impairment, bringing the signal amplitude back to the WT CTRL mice value (Fig. 4c). Subsequently, we explored if ABX-driven manipulation of the gut microbiota could improve behavioral impairments in CDD mice. First, we investigated and scored nest building (Fig. 4d), an instinctive/spontaneous behavior. Mice are highly motivated to build a nest as it has several functions related to survival and reproduction, including regulating body temperature, protecting from predators or aggressive cagemates, sheltering from cold or draft, and increasing litter survival (23). CDKL5 KO CTRL mice showed impaired nest building behavior compared to WT littermates; however, ABX treatment was unable to increase nest building scores in the mutants (Fig. 4e and Suppl. Fig. 5). Then, we scored hindlimb clasping (Fig. 4f), which is a signature of altered motor coordination and has been previously observed in CDKL5 KO mice (21). Unfortunately, due to the early test age, it was not possible to detect this motor deficit in CDKL5 KO CTRL mice, and ABX had no impact on this behavior (Fig. 4g).

Finally, we assessed exploration using the Y-maze (Fig. 4h). The CDKL5 KO CTRL mice performed a higher number of total entries than the WT CTRL mice (Fig. 4i) and displayed an increase in total distance moved and velocity (Fig. 4l,m). This hyperactive phenotype was normalized to the level of the WT littermates by manipulating the gut microbiota using ABX (Fig. 4i-m). In addition, CDKL5 KO CTRL mice performed worse than their WT littermates in the percentage of alternation in the three arms of the maze. Although the integrated statistics did not reveal significant results, the ABX tended to normalize this impairment in CDKL5 KO mice (Fig. 4n).

In summary, targeting gut microbiota dysbiosis in CDKL5 deficient mice was able to improve functional properties of cortical neurons, and ameliorated specific behavioral abnormalities.

### ABX-driven gut microbiota manipulation influence microglia phenotype in CDKL5 KO mice

The intestinal microbiota has emerged as a significant factor influencing the maturation, differentiation, and functionality of microglia cells (38,39). Interestingly, microglia from CDKL5 mutants displayed a phenotype typical of an activation state (40). Thus, based on this information, we explored whether ABX, which counteracts dysbiosis, could ameliorate the potential state of neuroinflammation in the CDKL5 KO brain by acting on microglia cells.

Three-dimensional morphological analysis of microglia cell bodies in the visual cortex (Fig. 5a) indicated no significant changes in soma area or soma volume in CDKL5 KO CTRL mice compared to WT mice (Fig. 5b-c). However, ABX treatment significantly decreased both soma area and volume with respect to WT and CDKL5 KO CTRL animals (Fig. 4b-c). In addition, CDKL5 KO CTRL mice displayed a significant reduction in microglia sphericity in comparison with WT littermates (Fig. 5d). Notably, this feature was rescued by ABX administration in CDKL5 KO mice (Fig. 5d). Since ameboid microglia lose the spherical shape of their cell body, a reduction in this characteristic might indicate that CDKL5 KO microglia are on the verge of activation in response to inflammatory signals, potentially emanating from a dysbiotic microbiota. administration of ABX may restore the cells to a surveillance state.

**FIG.5.**
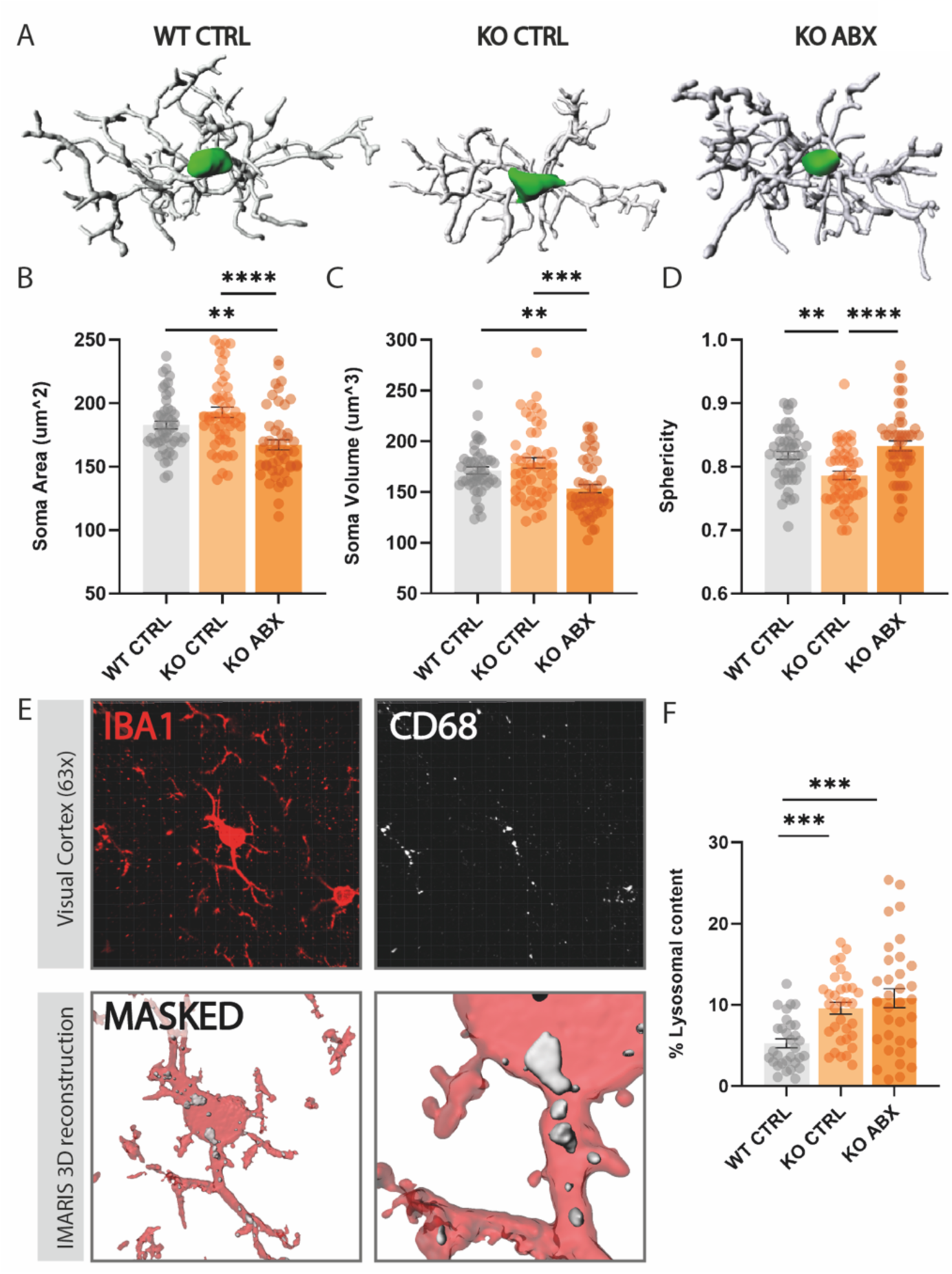
Microglia soma size analysis and expression of the lysosomal marker CD68 in CDKL5 KO mice visual cortex and effects of ABX administration. **(A)** Three-dimensional soma shape reconstruction of representative microglial cells from each experimental group. **(B)** Microglia soma area (um^2) (Kruskal-Wallis test p<0.0001, multiple comparisons Dunn’s post-hoc test; WT CTRL versus KO CTRL p=0.407, WT CTRL versus KO ABX p=0.007, KO CTRL versus KO ABX p<0.0001). **(C)** Microglia soma Volume (um^3) (Kruskal-Wallis test p=0.0002, multiple comparisons Dunn’s post-hoc test; WT CTRL versus KO CTRL p>0.999, WT CTRL versus KO ABX p=0.002, KO CTRL versus KO ABX p=0.0004). **(D)** Microglia sphericity (Kruskal-Wallis test p<0.0001, multiple comparisons Dunn’s post-hoc test; WT CTRL versus KO CTRL p=0.004, WT CTRL versus KO ABX p=0.698, KO CTRL versus KO ABX p<0.0001). **(E)** Representative volumes reconstruction of IBA1- and CD68-immunostained microglia. **(F)** % of Lysosomal content in IBA-1 immunostained microglial cells (Kruskal-Wallis test p<0.0001, multiple comparisons Dunn’s post-hoc test; WT CTRL versus KO CTRL p=0.0005, WT CTRL versus KO ABX p=0.0002, KO CTRL versus KO ABX p>0.999). Error bars represent SEM. Circles represent single cells.

To gain further insight on microglia function, we quantified their lysosomal content by CD68 staining (Fig. 5e). Our analysis revealed that CDKL5 KO microglia exhibited elevated expression of CD68 compared to WT mice, irrespective of the presence of an intact microbiota (Fig. 5f). This piece of data may suggest a higher phagocytic capacity of CDKL5 KO microglia with respect to WT animals.

Finally, microglia complexity (Fig. 6a) was investigated by quantifying filament length, number of branching points and number of terminal points. Intriguingly, CDKL5 KO microglia displayed a significant decrease in all of the above-mentioned parameters with respect to WT littermates, and ABX administration completely rescued the phenotype (Fig. 6b-d). Sholl analysis further demonstrated a reduction in CDKL5 KO CTRL cell complexity, due to a decrease of intersection numbers at specific soma distances (Suppl. Table 3), which was restored to WT levels by ABX treatment (Fig. 6e-g).

**FIG.6.**
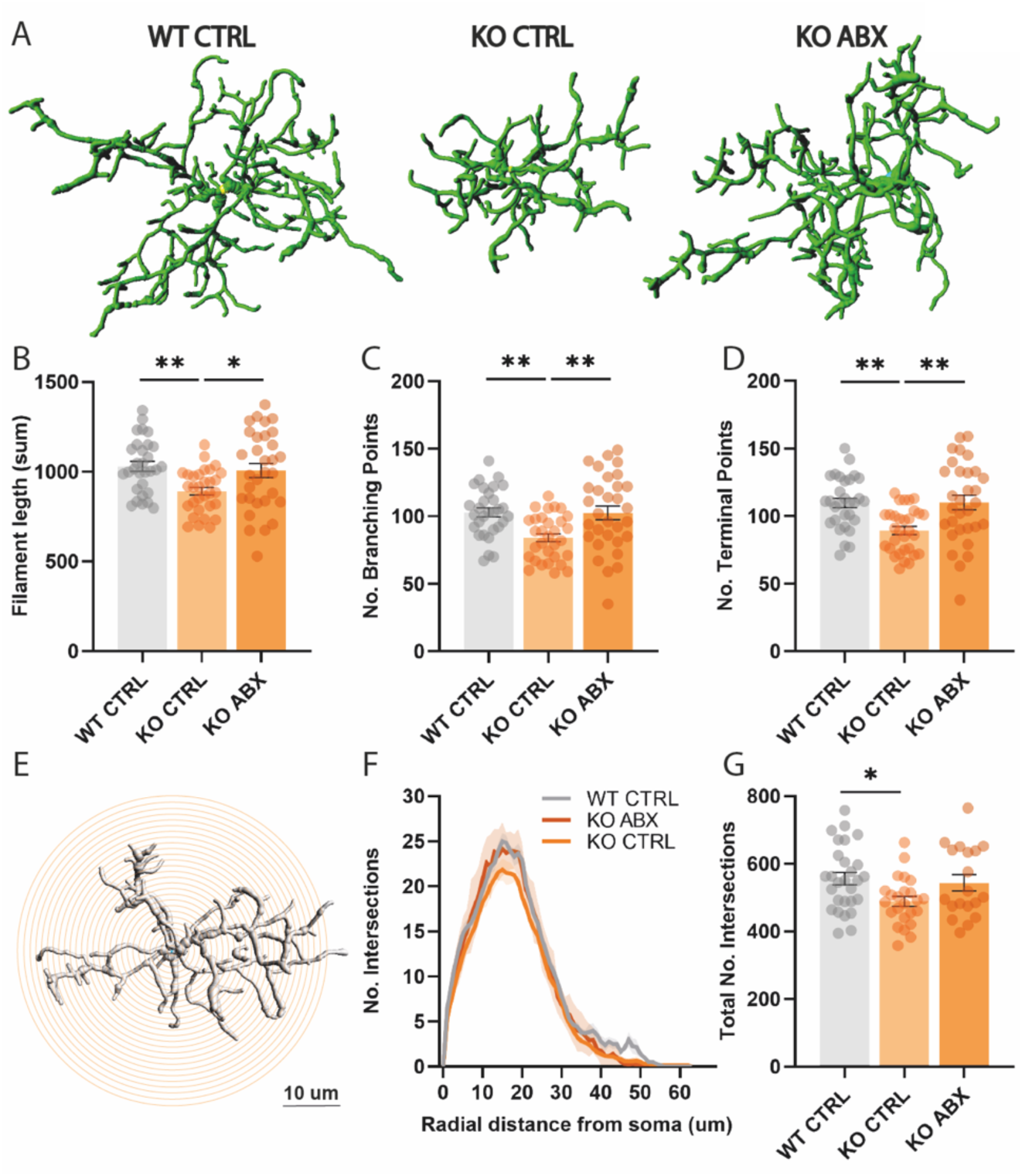
ABX treatment rescues microglia arborization and complexity. **(A)** Three-dimensional reconstruction of representative microglial cells arborization from each experimental group. **(B)** Filament length (sum) (Kruskal-Wallis test p=0.003, multiple comparisons Dunn’s post-hoc test; WT CTRL versus KO CTRL p=0.003, WT CTRL versus KO ABX p>0.999, KO CTRL versus KO ABX p=0.030). **(C)** Number of branching points (Kruskal-Wallis test p=0.0008, multiple comparisons Dunn’s post-hoc test; WT CTRL versus KO CTRL p=0.002, WT CTRL versus KO ABX p>0.999, KO CTRL versus KO ABX p=0.005). **(D)** Number of Terminal points (Kruskal-Wallis test p=0.0005, multiple comparisons Dunn’s post-hoc test; WT CTRL versus KO CTRL p=0.002, WT CTRL versus KO ABX p>0.999, KO CTRL versus KO ABX p=0.002). Circles represent single cells. **(E)** Representative image of a Sholl dendritic analysis in reconstructed IBA1-immunostained microglia. **(F)** Sholl analysis of microglial cells. (Two-way ANOVA interaction distance*experimental_group p=0.349, distance factor p<0.0001, experimental_group factor p<0.0001). Error bars represent SEM. Circles represent single cells.

Overall, our investigation of microglia phenotype suggests a potential state of activation in the CDKL5 KO cortex, that could be counteracted by manipulating the intestinal microbiota with ABX.

## DISCUSSION

Our investigation identified distinct variations in the developmental maturation pattern of the intestinal microbiota between two CDD mouse models and their WT littermates. As expected from the influence of various factors, including genetics, diet, and lifestyle, on the gut microbiota composition (41), the alterations present in CDD mice were affected by the specific model, age, and facility. However, we found a noteworthy CDD microbial signature characterized by dysregulated bacterial taxa that were consistently altered regardless of facility.

In both the Pisa and Berlin facilities, the major distinctions between CDKL5 KO animals and WT littermates were observed at juvenile ages. Alpha diversity was decreased at P25 and P32 in the Berlin vivarium, and a tendency for a decrease was observed in Pisa at the same ages. Similarly, in an Italian cohort of patients affected by CDD, fecal microbiota alpha diversity was diminished compared to healthy individuals (16). The Pisa CDKL5 KO mouse also displayed specific alterations in the gut microbiota: taxa belonging to the Phylum Proteobacteria were significantly enriched in these mice, at both P25 and P32 (i.e. orders Burkholderiales, Pasteurellales, families *Enterobacteriaceae*, *Alcaligenaceae*, genus *Sutterella*). These microbes were suggested as potential markers of microbiota instability and thus as a predisposition to disease onset (31). Blooming of Proteobacteria has been associated with various disease conditions, such as IBD, metabolic disorders and non-alcoholic fatty liver disease (NAFLD) (30). Indeed, the *Enterobacteriaceae* family is increased in type 2 diabetes patients (42), and the class Gammaproteobacteria was increased in children affected by NAFLD with respect to healthy controls (43). In addition, an enrichment of Proteobacteria has been found in mouse models of IBD (44–46), suggesting a potential state of intestinal inflammation in the CDD mouse model. Interestingly, recent reports have linked Proteobacteria to the modulation of the gut-brain axis. A significant increase in gut Proteobacteria associated with neuroinflammation, and accompained by exploration impairment and anxiety-like behavior was observed in a mouse model of diet induced obesity (47). Importantly, an enrichment in *Enterobacteriaceae* was observed in patients affected by CDD and manifesting severe GI disturbances compared to patients with mild to normal GI features (16), indicating similarities in the gut microbiota profile of mouse model and human subjects.

While the CDKL5 KO mice from the Berlin vivarium showed no differences at P32 compared to WT littermates, they showed enrichment in the Bacteroidaceae family at P25, which was also found in the Pisa CDKL5 KO mouse colony, demonstrating a common feature of the CDD microbiota. The expansion of *Bacteroidaceae* has been associated with colitis in mouse models (48,49), inflammation in cirrhotic patients (50), Alzheimer disease and IBD patients (51). Therefore, the blooming of *Bacteroidaceae* could be an early sign of dysbiosis in the CDD mouse. On the other hand, *Bifidobacterium longum* was underrepresented in the two CDD mouse models compared to their WT controls at P70. Multiple strains of *B. longum* have been formulated as probiotics due to their ability to elicit various health benefits. They have demonstrated efficacy in conditions such as IBD, cardiovascular diseases, respiratory illnesses, anxiety, depression and cognitive functions (52). Therefore, the reduction of *B. longum* in our CDD mouse models suggests a potential imbalance in the composition of gut microbes, characterized by a deficiency of beneficial bacteria that are crucial for maintaining intestinal homeostasis.

To further dissect additional shared taxa between the Pisa and Berlin CDD mouse models, we performed LEfSe analysis subtracting the variable “mouse facility”. This analysis highlighted the order Burkholderiales and Bacteroidales which are commonly enriched in the P25 CDKL5 KO mouse. In particular, *Bacteroides rodentium* was increased in CDKL5 KO at both P25 and P32, but no specific effects of this taxa on the gut-microbiota-brain axis have been reported so far. Among the common taxa decreased at P70, we found *Allobaculum. Allobaculum* is considered a beneficial microorganism especially in the context of hypertension, and it is negatively correlated with colonic expression of anti-inflammatory genes such as Foxp3 and IL-10 (53,54).

In summary, our multi-site experiment showed an interaction between mouse genetics, i.e. the absence of CDKL5 protein, and environmental conditions, i.e two different and geographically distant facilities, in shaping the intestinal microbiota. Previous studies suggest that the environment dominates over genetics (55) in determining the gut microbiota composition, and our results largely confirm this finding. As the DNA was extracted from all the fecal samples simultaneously and the sequencing was performed at the same time, we could exclude differences due to the sample processing. On the other hand, the standard chow used in the two facilities, the type of bedding and/or other environmental factors, could explain the dissimilarities between the Pisa and Berlin mice. Our efforts to identify microorganisms that are consistently altered in the CDKL5 KO mouse model revealed distinct taxa at specific ages. These taxa could be regarded as the CDKL5 KO signature microbes, suggesting a possible influence of genetics and indicating that the absence of functional CDKL5 might specifically shape the composition of the gut microbiota. The mechanisms by which impaired in CDKL5 functions could influence the maturation or growth of specific microorganisms is still unknown, and may suggest a role for CDKL5 in the enteric nervous system, or in the different types of cells of the intestine.

### Treating dysbiosis in CDKL5 KO mice improved phenotypical impairment

The presence of a pro-inflammatory dysbiotic microbial community in CDD mice raises the intriguing possibility that the intestinal microbiota of CDD mice may contribute to the symptoms in these mice. Our results showing that several symptoms are ameliorated by the administration of an antibiotic treatment to CDKL5 KO mice support this possibility. In particular, administration of ABX was able to improve visual response amplitude in CDKL5 KO mice. Visual alterations, also known as cortical visual impairment (CVI), were found to be present in 75% of CDD patients. CVI correlated with reduced achievement of milestones, and is a major feature of CDD that can impact on the quality of life and neurodevelopmental outcome of affected individuals (3,36). In particular, visual acuity, a quantifiable measure of visual function, has been found to be altered in CDD mice (56). The extent of impairment in visual acuity is linked to patients’ gross motor ability, suggesting that it can be used as an outcome measure in both pre-clinical and clinical studies of CDD (57). Improving this key symptom by targeting dysbiosis emphasizes the impact of abnormal gut-brain interaction in CDD beyond GI dysfunction.

A major challenge in gut-brain axis studies is to unveil the cellular underpinnings and potential mediators of the gut microbiota action on a distal tissue. Since subclinical inflammation is a characteristic feature of patients affected by CDD and a crosstalk between peripheral and central inflammatory processes has been proposed, we explored microglia cell behavior. Microglia, the predominant immune cells of the brain parenchyma, are key actors in neuroinflammation (58). Nevertheless, they are involved in synaptic remodeling and plasticity of neuronal circuits (59–64). Intriguingly, gut signals seem to influence microglia morphology and function (38,39,65,66), and in the CDKL5 KO mouse microglia cells are in a state of activation (40). We confirmed this result by observing a decrease in soma sphericity and an increase in lysosomal content in CDKL5 KO mice. Importantly, counteracting dysbiosis increased the cell-body sphericity in CDKL5 KO mice to the level of WT littermates. Finally, other microglia features linked to cellular structural complexity were found to be altered in CDKL5 KO mice. ABX treatment completely rescued this morphological phenotype, which could be linked to the abilitiy of microglia to sense and surveille the surrounding environment and thus prevent potential harmful pathogens/ molecules or maladaptive activation of the neuronal network.

### Translational perspectives in targeting the intestinal microbiota in CDD

Our study underscores the potential efficacy of targeting the gut microbiota to alleviate the symptoms of CDD. Employing a multisite approach, we identified a dysbiosis in CDD mouse models that mirrors the differences observed in patients. The presence of potentially pro-inflammatory taxa (e.g., *Bacteroidaceae*, *Enterobacteriaceae*) and a decrease in beneficial bacteria, such as *Bifidobacterium longum,* could contribute to an inflammatory state that activates microglial cells and impacts the severity of neurological symptoms. The ABX treatment improved neurological outcomes and microglia morphology, indicating partial relief from potential CDD-related inflammation. Notably, intestinal inflammation in humans has been linked to seizures and altered responsiveness to anti-seizure medication (67). This is a key aspect to consider in CDD patients who are affected by refractory epilepsy who have subclinical smoldering inflammation (6) and immune dysregulation (5). Our findings, therefore suggest harnessing the modifiable factor of gut microbiota as a novel target to alleviate symptom severity by mitigating peripheral and potentially brain low-grade inflammation. Easily applicable approaches such as probiotics tailored to the CDD fecal microbiota profile, prebiotics, and dietary supplements could offer almost side-effect-free interventions. Moreover, promising results have been obtained in ameliorating both neurological and GI symptoms in ASD patients via fecal transplantation (68–70), which, based on our investigation, could be conceivable for patients affected by CDD.

In conclusion, our pioneering exploration of the gut microbiota-brain axis in CDD unveils promising avenues for novel therapeutic strategies. We propose transformative treatments based on targeting the intestinal microbiota that hold the potential to alleviate symptoms and improve the overall well-being of patients with CDD.

## Acknowledgments

We thank all the members of Tognini’s team for insightful comments and feedback regarding the experiments. Special thanks to Valentino Totaro and Dr. Raffaele Mazziotti for their support with Python, and Dr. Leonardo Lupori for his assistance with the IOS technique. We also thank Giocchino Incandela, Francesca Biondi and Dr. Silvia Burchielli for their help in the mouse facility. This research was in part supported by Telethon Grant GSP21001 and by NextGenerationEU Italian Ministry of University and Research M4.C2-PNRR YOUNG MSCA_0000081 iNsPIReD to PT.

## Disclosures

The authors declare no competing interests.

16S rRNA-seq data will be publicly available at Zenodo upon acceptance of the manuscript.

## REFERENCES

1. Jakimiec M, Paprocka J, Śmigiel R (2020): CDKL5 Deficiency Disorder—A Complex Epileptic Encephalopathy. Brain Sciences, vol. 10. p 107.

2. Leonard H, Downs J, Benke TA, Swanson L, Olson H, Demarest S (2022): CDKL5 deficiency disorder: clinical features, diagnosis, and management. Lancet Neurol 21: 563–576.

3. Demarest ST, Olson HE, Moss A, Pestana-Knight E, Zhang X, Parikh S, et al. (2019): CDKL5 deficiency disorder: Relationship between genotype, epilepsy, cortical visual impairment, and development. Epilepsia 60: 1733–1742.

4. Mangatt M, Wong K, Anderson B, Epstein A, Hodgetts S, Leonard H, Downs J (2016): Prevalence and onset of comorbidities in the CDKL5 disorder differ from Rett syndrome. Orphanet J Rare Dis 11: 39.

5. Leoncini S, De Felice C, Signorini C, Zollo G, Cortelazzo A, Durand T, et al. (2015) : Cytokine Dysregulation in MECP2- and CDKL5-Related Rett Syndrome: Relationships with Aberrant Redox Homeostasis, Inflammation, and ω-3 PUFAs. Oxid Med Cell Longev 2015: 421624.

6. Cortelazzo A, de Felice C, Leoncini S, Signorini C, Guerranti R, Leoncini R, et al. (2017): Inflammatory protein response in CDKL5-Rett syndrome: evidence of a subclinical smouldering inflammation. Inflamm Res 66: 269–280.

7. Saurman V, Margolis KG, Luna RA (2020): Autism Spectrum Disorder as a Brain-Gut-Microbiome Axis Disorder. Dig Dis Sci 65: 818–828.

8. McElhanon BO, McCracken C, Karpen S, Sharp WG (2014): Gastrointestinal symptoms in autism spectrum disorder: a meta-analysis. Pediatrics 133: 872–883.

9. Doshi-Velez F, Ge Y, Kohane I (2014): Comorbidity clusters in autism spectrum disorders: an electronic health record time-series analysis. Pediatrics 133: e54–63.

10. Richdale AL, Schreck KA (2009): Sleep problems in autism spectrum disorders: prevalence, nature, & possible biopsychosocial aetiologies. Sleep Med Rev 13: 403– 411.

11. Damiani F, Cornuti S, Tognini P (2023): The gut-brain connection: Exploring the influence of the gut microbiota on neuroplasticity and neurodevelopmental disorders. Neuropharmacology 231: 109491.

12. Murakami M, Tognini P (2019): The Circadian Clock as an Essential Molecular Link Between Host Physiology and Microorganisms. Front Cell Infect Microbiol 9: 469.

13. Ding X, Xu Y, Zhang X, Zhang L, Duan G, Song C, et al. (2020): Gut microbiota changes in patients with autism spectrum disorders. J Psychiatr Res 129: 149–159.

14. Borghi E, Borgo F, Severgnini M, Savini MN, Casiraghi MC, Vignoli A (2017): Rett Syndrome: A Focus on Gut Microbiota. Int J Mol Sci 18. 10.3390/ijms18020344

15. Strati F, Cavalieri D, Albanese D, De Felice C, Donati C, Hayek J, et al. (2016): Altered gut microbiota in Rett syndrome. Microbiome 4: 41.

16. Borghi E, Xynomilakis O, Ottaviano E, Ceccarani C, Viganò I, Tognini P, Vignoli A (2023, December 3): Gut microbiota profile in CDKL5 deficiency disorder patients as a potential marker of clinical severity. bioRxiv. 10.1101/2023.12.01.569361

17. Mitrea L, Nemeş S-A, Szabo K, Teleky B-E, Vodnar D-C (2022): Guts Imbalance Imbalances the Brain: A Review of Gut Microbiota Association With Neurological and Psychiatric Disorders. Front Med 9: 813204.

18. Sultan S, El-Mowafy M, Elgaml A, Ahmed TAE, Hassan H, Mottawea W (2021): Metabolic Influences of Gut Microbiota Dysbiosis on Inflammatory Bowel Disease. Front Physiol 12: 715506.

19. Yu LC-H (2018): Microbiota dysbiosis and barrier dysfunction in inflammatory bowel disease and colorectal cancers: exploring a common ground hypothesis. J Biomed Sci 25: 1–14.

20. Hamjane N, Mechita MB, Nourouti NG, Barakat A (2023): Gut microbiota dysbiosis - associated obesity and its involvement in cardiovascular diseases and type 2 diabetes. A systematic review. Microvasc Res 151: 104601.

21. Amendola E, Zhan Y, Mattucci C, Castroflorio E, Calcagno E, Fuchs C, et al. (2014): Mapping pathological phenotypes in a mouse model of CDKL5 disorder. PLoS One 9: e91613.

22. Zhu J-W, Li Y-F, Wang Z-T, Jia W-Q, Xu R-X (2016): Toll-Like Receptor 4 Deficiency Impairs Motor Coordination. Front Neurosci 10: 33.

23. Deacon RMJ (2006): Assessing nest building in mice. Nat Protoc 1: 1117–1119.

24. Thukral AK (2017): A review on measurement of Alpha diversity in biology. Agricultural Research Journal, vol. 54. p 1.

25. Pinart M, Dötsch A, Schlicht K, Laudes M, Bouwman J, Forslund SK, et al. (2021): Gut Microbiome Composition in Obese and Non-Obese Persons: A Systematic Review and Meta-Analysis. Nutrients 14. 10.3390/nu14010012

26. Hirano A, Umeno J, Okamoto Y, Shibata H, Ogura Y, Moriyama T, et al. (2018): Comparison of the microbial community structure between inflamed and non-inflamed sites in patients with ulcerative colitis. J Gastroenterol Hepatol. 10.1111/jgh.14129

27. Li Z, Zhou J, Liang H, Ye L, Lan L, Lu F, et al. (2022): Differences in Alpha Diversity of Gut Microbiota in Neurological Diseases. Front Neurosci 16: 879318.

28. Bray JR, Curtis JT (1957): An Ordination of the Upland Forest Communities of Southern Wisconsin.

29. Segata N, Izard J, Waldron L, Gevers D, Miropolsky L, Garrett WS, Huttenhower C (2011): Metagenomic biomarker discovery and explanation. Genome Biol 12: R60.

30. Rizzatti G, Lopetuso LR, Gibiino G, Binda C, Gasbarrini A (2017): Proteobacteria: A Common Factor in Human Diseases. Biomed Res Int 2017. 10.1155/2017/9351507

31. Shin N-R, Whon TW, Bae J-W (2015): Proteobacteria: microbial signature of dysbiosis in gut microbiota. Trends Biotechnol 33: 496–503.

32. Anand N, Gorantla VR, Chidambaram SB (2022): The Role of Gut Dysbiosis in the Pathophysiology of Neuropsychiatric Disorders. Cells 12. 10.3390/cells12010054

33. Morais LH, Schreiber HL, Mazmanian SK (2020): The gut microbiota–brain axis in behaviour and brain disorders. Nat Rev Microbiol 19: 241–255.

34. Mazziotti R, Lupori L, Sagona G, Gennaro M, Della Sala G, Putignano E, Pizzorusso T (2017): Searching for biomarkers of CDKL5 disorder: early-onset visual impairment in CDKL5 mutant mice. Hum Mol Genet 26: 2290–2298.

35. Lupori L, Sagona G, Fuchs C, Mazziotti R, Stefanov A, Putignano E, et al. (2019): Site-specific abnormalities in the visual system of a mouse model of CDKL5 deficiency disorder. Human Molecular Genetics, vol. 28. pp 2851–2861.

36. Brock D, Fidell A, Thomas J, Juarez-Colunga E, Benke TA, Demarest S (2021): Cerebral Visual Impairment in CDKL5 Deficiency Disorder Correlates With Developmental Achievement. J Child Neurol 36: 974–980.

37. Quintiliani M, Ricci D, Petrianni M, Leone S, Orazi L, Amore F, et al. (2021): Cortical Visual Impairment in CDKL5 Deficiency Disorder. Front Neurol 12: 805745.

38. Erny D, Hrabě de Angelis AL, Jaitin D, Wieghofer P, Staszewski O, David E, et al. (2015): Host microbiota constantly control maturation and function of microglia in the CNS. Nat Neurosci 18: 965–977.

39. Erny D, Dokalis N, Mezö C, Castoldi A, Mossad O, Staszewski O, et al. (2021): Microbiota-derived acetate enables the metabolic fitness of the brain innate immune system during health and disease. Cell Metab 33: 2260–2276.e7.

40. Galvani G, Mottolese N, Gennaccaro L, Loi M, Medici G, Tassinari M, et al. (2021): Inhibition of microglia overactivation restores neuronal survival in a mouse model of CDKL5 deficiency disorder. J Neuroinflammation 18: 155.

41. Parizadeh M, Arrieta M-C (2023): The global human gut microbiome: genes, lifestyles, and diet. Trends Mol Med 29: 789–801.

42. American Diabetes Association (2014): Diagnosis and classification of diabetes mellitus. Diabetes Care 37 Suppl 1: S81–90.

43. Michail S, Lin M, Frey MR, Fanter R, Paliy O, Hilbush B, Reo NV (2015): Altered gut microbial energy and metabolism in children with non-alcoholic fatty liver disease. FEMS Microbiol Ecol 91: 1–9.

44. Carvalho FA, Koren O, Goodrich JK, Johansson MEV, Nalbantoglu I, Aitken JD, et al. (2012): Transient inability to manage proteobacteria promotes chronic gut inflammation in TLR5-deficient mice. Cell Host Microbe 12: 139–152.

45. Selvanantham T, Lin Q, Guo CX, Surendra A, Fieve S, Escalante NK, et al. (2016): NKT Cell-Deficient Mice Harbor an Altered Microbiota That Fuels Intestinal Inflammation during Chemically Induced Colitis. J Immunol 197: 4464–4472.

46. Maharshak N, Packey CD, Ellermann M, Manick S, Siddle JP, Huh EY, et al. (2013): Altered enteric microbiota ecology in interleukin 10-deficient mice during development and progression of intestinal inflammation. Gut Microbes 4: 316–324.

47. Jeong M-Y, Jang H-M, Kim D-H (2019): High-fat diet causes psychiatric disorders in mice by increasing Proteobacteria population. Neurosci Lett 698: 51–57.

48. Harrison CA, Laubitz D, Ohland CL, Midura-Kiela MT, Patil K, Besselsen DG, et al. (2018): Microbial dysbiosis associated with impaired intestinal Na/H exchange accelerates and exacerbates colitis in ex-germ free mice. Mucosal Immunol 11: 1329–1341.

49. Ariake K, Ohkusa T, Sakurazawa T, Kumagai J, Eishi Y, Hoshi S, Yajima T (2000): Roles of mucosal bacteria and succinic acid in colitis caused by dextran sulfate sodium in mice. J Med Dent Sci 47: 233–241.

50. Piñero F, Vazquez M, Baré P, Rohr C, Mendizabal M, Sciara M, et al. (2019): A different gut microbiome linked to inflammation found in cirrhotic patients with and without hepatocellular carcinoma. Ann Hepatol 18: 480–487.

51. Wang D, Zhang X, Du H (2022): Inflammatory bowel disease: A potential pathogenic factor of Alzheimer’s disease. Prog Neuropsychopharmacol Biol Psychiatry 119: 110610.

52. Mills S, Yang B, Smith GJ, Stanton C, Ross RP (2023): Efficacy of alone or in multi-strain probiotic formulations during early life and beyond. Gut Microbes 15: 2186098.

53. Dan X, Mushi Z, Baili W, Han L, Enqi W, Huanhu Z, Shuchun L (2019): Differential Analysis of Hypertension-Associated Intestinal Microbiota. Int J Med Sci 16: 872– 881.

54. Richards EM, Pepine CJ, Raizada MK, Kim S (2017): The Gut, Its Microbiome, and Hypertension. Curr Hypertens Rep 19: 36.

55. Rothschild D, Weissbrod O, Barkan E, Kurilshikov A, Korem T, Zeevi D, et al. (2018): Environment dominates over host genetics in shaping human gut microbiota. Nature 555: 210–215.

56. Mazziotti R, Lupori L, Sagona G, Gennaro M, Della Sala G, Putignano E, Pizzorusso T (2017): Searching for biomarkers of CDKL5 disorder: early-onset visual impairment in CDKL5 mutant mice. Hum Mol Genet 26: 2290–2298.

57. Olson HE, Costantini JG, Swanson LC, Kaufmann WE, Benke TA, Fulton AB, et al. (2021): Cerebral visual impairment in CDKL5 deficiency disorder: vision as an outcome measure. Dev Med Child Neurol 63: 1308–1315.

58. Leng F, Edison P (2020): Neuroinflammation and microglial activation in Alzheimer disease: where do we go from here? Nat Rev Neurol 17: 157–172.

59. Microglial Remodeling of the Extracellular Matrix Promotes Synapse Plasticity (2020): Cell 182: 388–403.e15.

60. Sipe GO, Lowery RL, Tremblay M-È, Kelly EA, Lamantia CE, Majewska AK (2016): Microglial P2Y12 is necessary for synaptic plasticity in mouse visual cortex. Nat Commun 7: 1–15.

61. Miyamoto A, Wake H, Ishikawa AW, Eto K, Shibata K, Murakoshi H, et al. (2016): Microglia contact induces synapse formation in developing somatosensory cortex. Nat Commun 7: 12540.

62. Parkhurst CN, Yang G, Ninan I, Savas JN, Yates JR 3rd, Lafaille JJ, et al. (2013): Microglia promote learning-dependent synapse formation through brain-derived neurotrophic factor. Cell 155: 1596–1609.

63. Hashimoto A, Kawamura N, Tarusawa E, Takeda I, Aoyama Y, Ohno N, et al. (2023): Microglia enable cross-modal plasticity by removing inhibitory synapses. Cell Rep 42: 112383.

64. Eichler A, Kleidonas D, Turi Z, Fliegauf M, Kirsch M, Pfeifer D, et al. (2023): Microglial Cytokines Mediate Plasticity Induced by 10 Hz Repetitive Magnetic Stimulation. J Neurosci 43: 3042–3060.

65. Huang Y, Wu J, Zhang H, Li Y, Wen L, Tan X, et al. (2023): The gut microbiome modulates the transformation of microglial subtypes. Mol Psychiatry 28: 1611–1621.

66. Lupori L, Cornuti S, Mazziotti R, Borghi E, Ottaviano E, Cas MD, et al. (2022): The gut microbiota of environmentally enriched mice regulates visual cortical plasticity. Cell Rep 38: 110212.

67. De Caro C, Leo A, Nesci V, Ghelardini C, di Cesare Mannelli L, Striano P, et al. (2019): Intestinal inflammation increases convulsant activity and reduces antiepileptic drug efficacy in a mouse model of epilepsy. Sci Rep 9: 13983.

68. Kang D-W, Adams JB, Gregory AC, Borody T, Chittick L, Fasano A, et al. (2017): Microbiota Transfer Therapy alters gut ecosystem and improves gastrointestinal and autism symptoms: an open-label study. Microbiome 5: 10.

69. Kang D-W, Adams JB, Coleman DM, Pollard EL, Maldonado J, McDonough-Means S, et al. (2019): Long-term benefit of Microbiota Transfer Therapy on autism symptoms and gut microbiota. Sci Rep 9: 5821.

70. Li N, Chen H, Cheng Y, Xu F, Ruan G, Ying S, et al. (2021): Fecal Microbiota Transplantation Relieves Gastrointestinal and Autism Symptoms by Improving the Gut Microbiota in an Open-Label Study. Front Cell Infect Microbiol 11: 759435.

