## Supplementary methods and Figures for "Multi-Site Investigation of Gut Microbiota in CDKL5 Deficiency Disorder Mouse Models: Targeting Dysbiosis to Improve Neurological Outcomes"

### SUPPLEMENTS

#### SUPPLEMENTARY METHODS

##### 16S rRNA-sequencing method details

Library preparation and sequencing. Libraries were prepared by following Illumina 16S Metagenomic Sequencing Library Preparation protocol in two amplification steps: an initial 35 cycle PCR amplification using locus-specific PCR primers and a subsequent amplification that integrates relevant flow-cell binding domains and unique indices (NexteraXT Index Kit, FC-131- 1001/FC-131-1002).

Primer sequences used for amplification:

16S F (341F) 5'- CCTACGGGNGGCWGCAG -3'

16S R (805R) 5'- GGACTACHVGGGTATCTAATCC -3'

Libraries were sequenced on a MiSeq instrument (Illumina) in paired end 300-bp mode read length.

Bioinformatics analysis. Reads were de-multiplexed based on the Illumina indexing system. Where amplicon length was permissive with the respective sequencing length, 3'-ends of pairs were overlapped to generate consensus pseudo-reads, while the remainders were maintained as separated pairs. After, a clipping routine is applied to remove low-quality bases at 3'tails. Reads were then retained if they maintained a minimum length of 200 bp. Any primer sequence at 5'-ends was removed and not accounted for during the process.

Following the QIIME pipelines, the USEARCH algorithm (version 8.1.1756, 32-bit) allows the following steps: chimera filtering, grouping of replicate sequences, sorting sequences per decreasing abundance and operational taxonomic unit (OTU) identification. The OTU picking aims to group query sequences into clusters, represented by centroids. Each centroid shares a level of similarity with their member sequences.

All reads were used in the analysis if they maintain a minimum length of 200 bp after removal of primer sequence and low quality bases. Paired reads with permissive overlap at their 3'-ends were merged to single fragment and used as such to improve assignment accuracy. Reads that did not support overlap were maintained in the pool for downstream processing. Sequence clustering was performed using the open-reference method with a threshold set at 97%. Sequences that passed a pre-filter step, ensuring a minimum identity of 90% with any sequence in the reference database, contributed to OTU formation. The "open-reference" analysis generated OTUs, each requiring a minimum of 2 sequenced fragments.

Rarefaction curves end-points and normalization of counts for diversity analysis are set to 50% of the target sequencing coverage (i.e. for 100,000 fragments a cutoff of 50,000 fragments is applied). The cutoff could be modified according to the sequencing yields. The total count was retained for taxonomic abundances estimation and used accordingly for ad-hoc statistical testing of taxonomic abundance when enquired.

The RDP classifier and Reference database were used to assign taxonomy with a minimum confidence threshold of 0.50. IGA Tech Reference database: 16S: modified GreenGene database (version 2013_8).

**Immunofluorescence analysis**

Mice were anesthetized with chloral hydrate (20ml/Kg BW) and perfused via intracardiac infusion with PBS and then 4% paraformaldehyde (PFA, w/vol, dissolved in 0.1 M phosphate buffer, pH 7.4). Brains were quickly removed and post-fixed overnight in PFA at 4 °C, then transferred to 30% sucrose (w/vol) solution. 45 µm coronal sections were cut on a freezing microtome (Leica) and free-floating sections were processed for immunofluorescence.

The cortical sections were incubated for 1 h in a blocking solution containing 5% BSA (w/vol) and 0.3% Triton X-100 (vol/vol) in PBS, and incubated overnight at 4 °C with anti-Iba-1 (cat. no. 019-19741, Wako) diluted 1:500, and anti-CD68 (cat. n MCA1957GA) 1:250, in PBS with 1% BSA (w/vol) and 0.1% Triton X-100 (vol/vol).

Sections were then washed with PBS and incubated for 2h at 22-24 °C with Alexa Fluor 568–conjugated secondary antibody (cat. no A11011, Invitrogen) and Alexa Fluor 647–conjugated secondary antibody (cat. no A21247, Invitrogen), which was added at a dilution of 1:500 in the same solution as the primary antibody.

Sections were washed three times with PBS and mounted on slides, then they were air-dried and coverslipped with Vectashield mounting medium (cat. H-1000,Vector Laboratories).

Imaging was performed on an LSM 900 confocal microscope (Zeiss, Oberkochen, Germany) using a Plan-Apochromat 63x, NA:1.4 oil objective.

The area of the visual cortex was defined based on the mouse brain atlas (Paxinos and Franklin's the Mouse Brain in Stereotaxic Coordinates).

The 3D reconstruction of microglial cells was performed on Z-stacks of ~ 40 μm, acquired with a z-step of 0.50 μm (for a final voxel size of 0.099 × 0.099 × 0.5 μm). Images were then processed for filament and soma reconstruction using respectively the “Filaments” and “Surfaces” functions of IMARIS software (Bitplane). Between 5 and 10 cortical cells were reconstructed per mouse. The analyzed morphological features were selected based on their relevance as indicators of microglia activation: filament length, number of branching points, number of terminal points, soma area, soma volume and soma sphericity (roundness) (1,2).

The phagocytic activity of microglial cells through CD68 quantitation was assessed on Z-stacks of 19.8 μm, acquired with a z-step of 1.98 μm (for each ROI, 10 fields of view containing phagocytes were acquired). The protocol used for the surface reconstruction of microglial cells and phagocytes was adapted from (3).

**Intrinsic Optical Imaging (IOS)**

Surgery. Surgery for IOS imaging was performed as described in (4). P30 CDKL5 KO and WT littermates were anesthetized with isoflurane (3% induction; 1% maintenance) and head fixed on a stereotaxic frame using ear bars. Body temperature was monitored using a heating pad and a rectal probe to maintain the animals’ body at 37ºC. A subcutaneous injection of lidocaine (2%) was provided to anesthetize the local area and the eyes were protected with a dexamethasone-based ointment (Tobradex, Alcon Novartis). The scalp was removed and the skull cleaned with saline. The skin was secured to the skull using cyanoacrylate and a thin layer of cyanoacrylate was poured over the exposed skull to attach a custom-made metal ring (9 mm internal diameter) centered over the binocular visual cortex. A thin layer of clear nail polish was applied over the area to ameliorate optical access. After surgery, the animals were allowed to recover fully in a heated box and monitored to ensure the absence of any sign of discomfort. Before any other experimental procedure, mice were left to recover for at least 4 days.

Imaging and data analysis. Mice were anesthetized with isoflurane (3% induction; 1% maintenance) and chlorprothixene anesthesia (1.5 mg/kg, i.p.). Images were visualized using a custom Leica microscope (Leica Microsystems). Red light illumination was provided by 6 individually addressable LEDs (WS2812) attached to the objective (Leica Z6 APO coupled with a Leica PlanApo 2.0X 10447178) by a custom 3D-printed conical holder. Visual stimuli were generated using Matlab Psychtoolbox and presented on a gamma-corrected 24” monitor (C24F390FHU).

Horizontal sine-wave gratings were presented in the binocular portion of the visual field enclosed in a Gaussian envelope spanning -10 to +10 degrees of azimuth and -5 to +60 (full monitor height) degrees of altitude, with a spatial frequency of 0.03 cycles per degree, mean luminance 20 cd/$m^2$ and a contrast of 90%. The stimulus consisted of the abrupt contrast reversal of a grating with a temporal frequency of 4 Hz for 1 second, time-locked with a 12-bit depth acquisition camera (PCO edge 5.5) using a parallel port trigger. The interstimulus time was 13 seconds. Frames were acquired at 30 fps with a resolution of 540 x 640 pixels. The signal was averaged for at least 8 groups of 20 trials, stimulating the contralateral eye to the recorded visual cortex. The signal was then downsampled in time to 10 fps and in space to 270 x 320 pixels. Fluctuations of reflectance (R) for each pixel were computed as the normalized difference from the average baseline ($\Delta R/R$). For each recording, an image representing the mean evoked response was computed by averaging frames between 0.5 to 2.5 seconds after stimulation. The mean image was then low-pass filtered with a 2D average square spatial filter (7 pixels). To select the binocular portion of the V1 for further analysis, a region of interest (ROI) was automatically calculated on the mean image of the response by selecting the pixels in the lowest 20% $\Delta R/R$ of the range between the maximal and minimal intensity pixel (5). To weaken background fluctuations a manually selected polygonal region of reference (ROR) was subtracted. The ROR was placed where no clear response, blood vessel artifact or irregularities of the skull were observed (6) . Mean evoked responses were quantitatively estimated as the average intensity inside the ROI.

**Detailed Statistical Analysis**

Gut microbiota analysis: to test whether two or more groups of samples were significantly different, PERMANOVA and principal coordinate analysis (PCoA) were calculated using the Python library scikit-bio (http://scikit-bio.org/) with, respectively, the skbio.stats.distance.permanova and skbio.stats.ordination.pcoa functions. Alpha diversity significance was calculated using Mann-Whitney statistics.

IOS experiments: Differences between groups were tested for significance using one way ANOVA. Tukey's multiple comparisons post hoc tests were performed, to correct for multiple hypothesis testing.

Behavioral analysis: Differences between groups on the Y-maze task were tested for significance using one way ANOVA. Nesting and clasping scores were analyzed using non-parametric Kruskal-Wallis test since normality and homoscedasticity assumptions were not respected. Tukey's multiple comparisons post hoc tests were performed, when appropriate, to correct for multiple hypothesis testing.

Microglia morphology: For the analysis of soma area, volume and sphericity, filament length, number of branching points and dendrite terminals, statistical differences between groups were assessed using non-parametric Kruskal-Wallis test, followed by Dunn's post hoc multiple comparisons test. For the Sholl analysis, a Two-Way ANOVA was applied for assessing interaction effects between distance from the soma and treatment, followed by Tukey's post hoc multiple comparisons test.

#### SUPPLEMENTARY TABLE DESCRIPTION

**Suppl. Table 1**: Relative abundance of bacteria in the fecal microbiota of mice belonging to Pisa Vivarium.

**Suppl. Table 2**: Relative abundance of bacteria in the fecal microbiota of mice belonging to Berlin Vivarium.

**Suppl. Table 3:** Two-way ANOVA Multiple Comparisons tests of Sholl Intersection analysis (referred to Figure 6f).

#### REFERENCES


1. Jurga AM, Paleczna M, Kuter KZ (2020): Overview of General and Discriminating Markers of Differential Microglia Phenotypes. *Front Cell Neurosci* 14: 198.
2. Franco-Bocanegra DK, Gourari Y, McAuley C, Chatelet DS, Johnston DA, Nicoll JAR, Boche D (2021): Microglial morphology in Alzheimer's disease and after Aβ immunotherapy. *Sci Rep* 11: 15955.
3. Schafer DP, Lehrman EK, Heller CT, Stevens B (2014): An engulfment assay: a protocol to assess interactions between CNS phagocytes and neurons. *J Vis Exp*. https://doi.org/10.3791/51482
4. Mazziotti R, Lupori L, Sagona G, Gennaro M, Della Sala G, Putignano E, Pizzorusso T (2017): Searching for biomarkers of CDKL5 disorder: early-onset visual impairment in CDKL5 mutant mice. *Hum Mol Genet* 26: 2290–2298.
5. Cang J, Kalatsky VA, Löwel S, Stryker MP (2005): Optical imaging of the intrinsic signal as a measure of cortical plasticity in the mouse. *Vis Neurosci* 22: 685–691.
6. Heimel JA, Hartman RJ, Hermans JM, Levelt CN (2007): Screening mouse vision with intrinsic signal optical imaging. *Eur J Neurosci* 25: 795–804.

#### SUPPLEMENTARY FIGURES

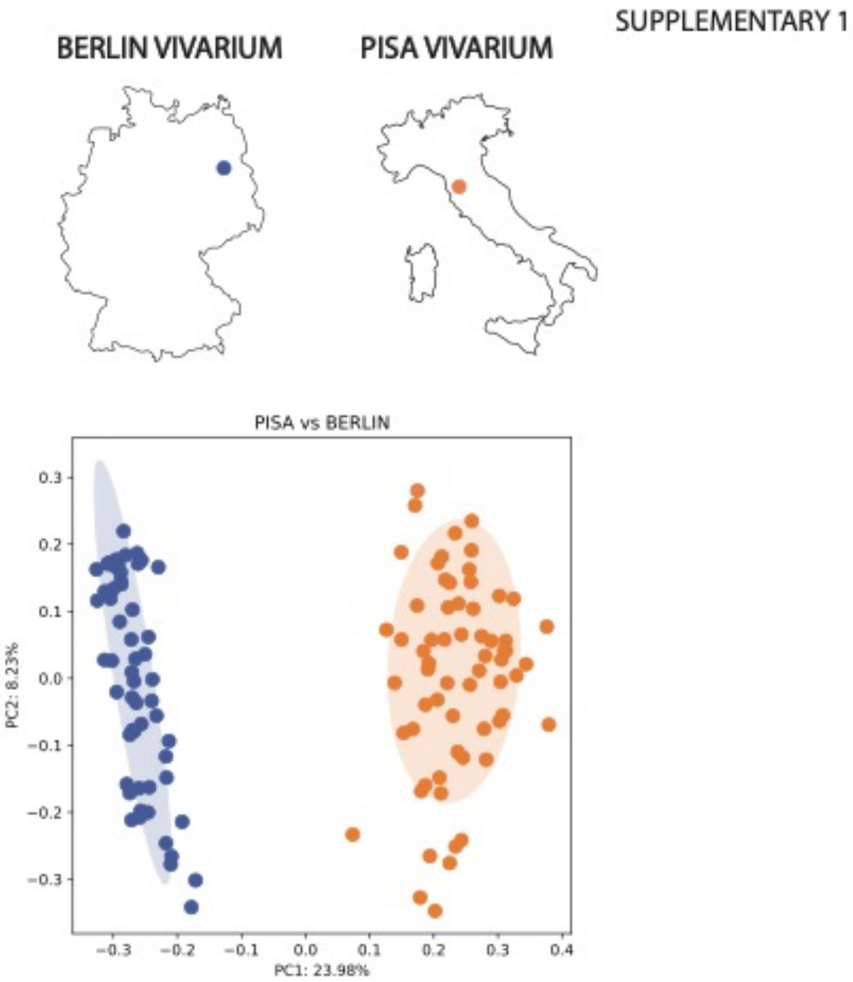


**SUPPL. Fig. 1** PcoA plot based on Bray-Curtis dissimilarity matrix showing beta diversity of all the WT and KO mice from Pisa (orange) and Berlin (blue) vivaria. The ellipses represent 95% confidence intervals for each group. Axes in the PCoA display the percentage of variation explained using Bray-Curtis dissimilarity.

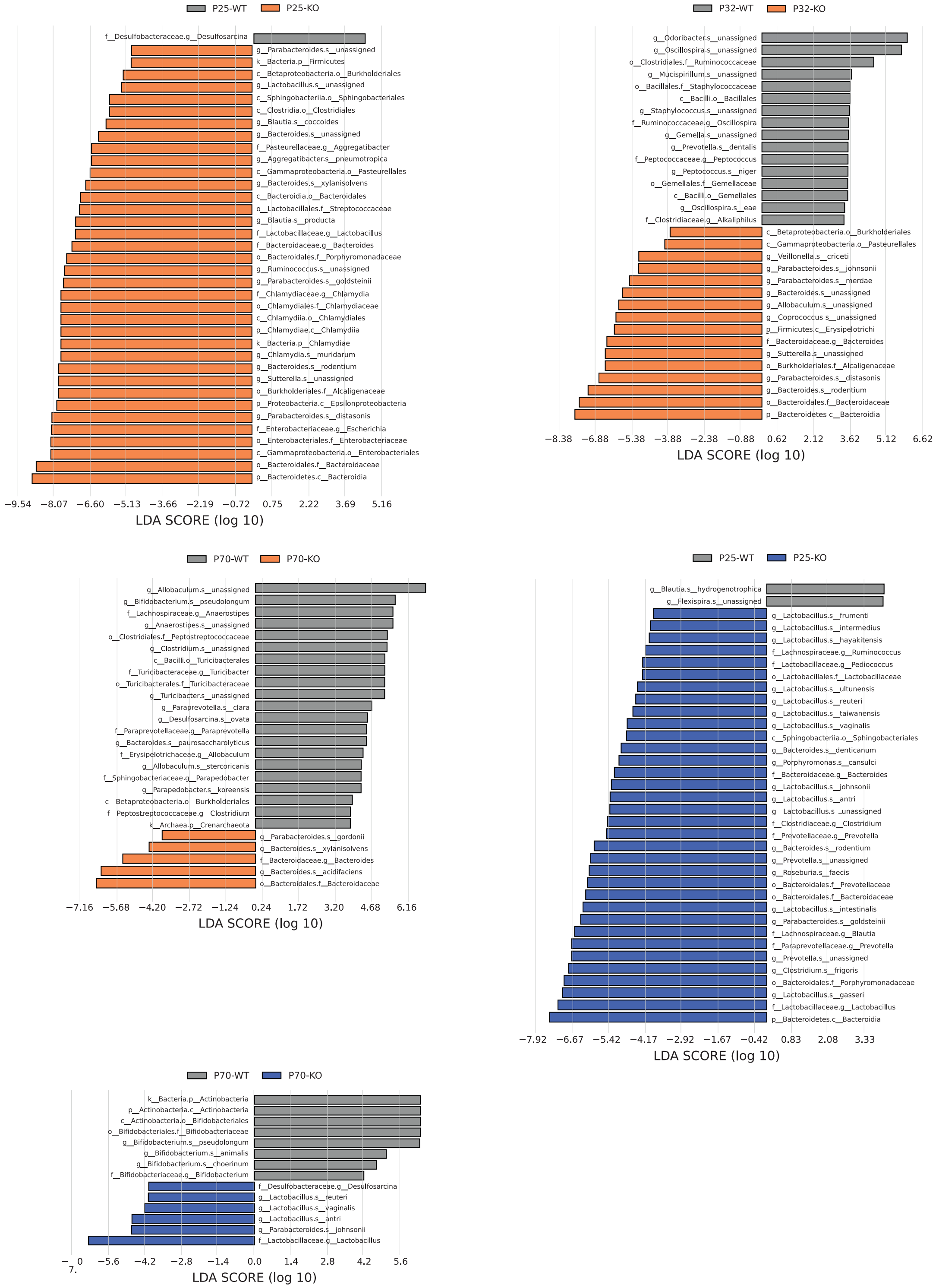


**SUPPL. Fig. 2.** Histogram of LDA scores (>2.5) obtained for bacterial taxa differentially abundant between WT (gray) and KO (orange for Pisa, blue for Berlin) mice at P25, P32 and P70.

SUPPLEMENTARY 3

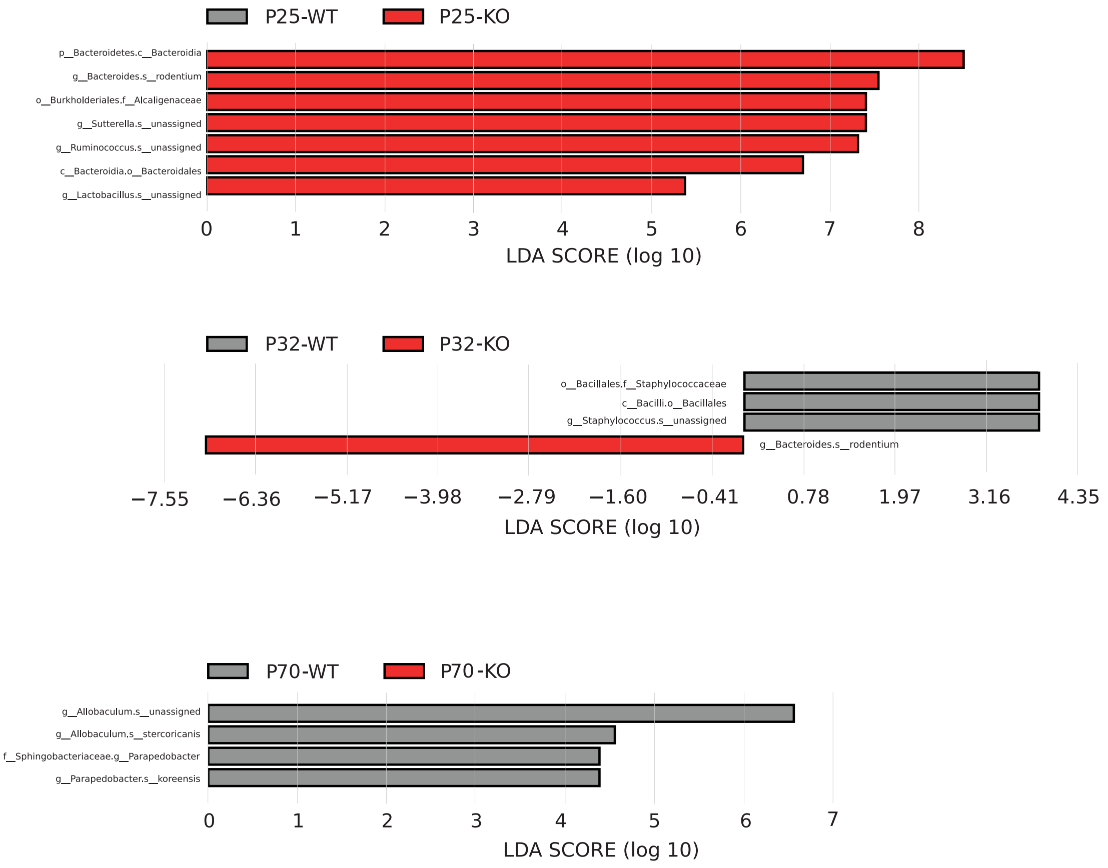


**SUPPL. Fig. 3.** Histogram of LDA scores (>2.5) obtained for bacterial taxa differentially abundant between WT (gray) and KO (red) mice subtracting the variable mouse facility, at P25, P32 and P70.

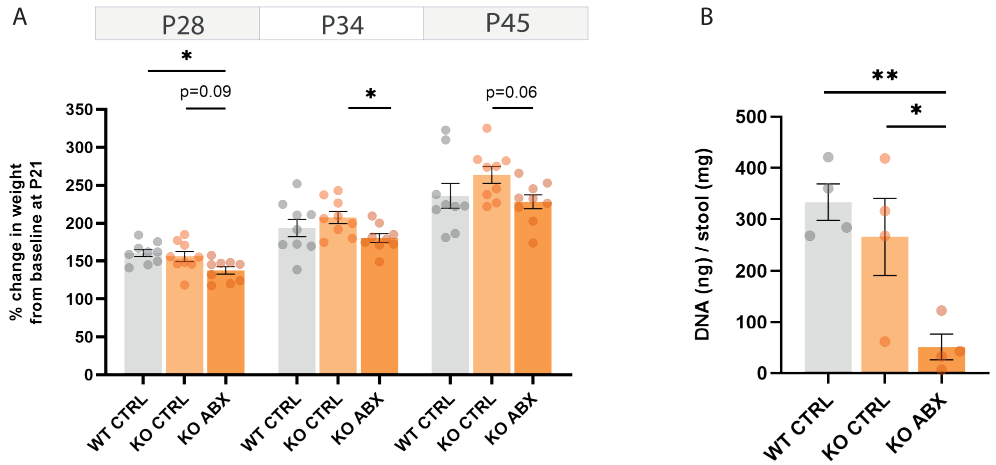


**SUPPL. Fig. 4.** Effects of the ABX treatment on body weight gain and fecal concentration of bacterial DNA. **(A)** % of change in body weight from the baseline (P21) at the time point of P28, P34 and P45. Error bars represent SEM. Circles represent single experimental subjects. (RM Two-way ANOVA interaction time*experimental_group p=0.274, time factor p<0.0001, experimental_group factor p=0.053, subject factor p=0.002; multiple comparisons Tukey post-hoc test (simple effects within the time factor); P28 WT CTRL versus KO CTRL p=0.832, P28 WT CTRL versus KO ABX p=0.008, P28 KO CTRL versus KO ABX p=0.096, P34 WT CTRL versus KO CTRL p=0.589, P34 WT CTRL versus KO ABX p=0.570, P34 KO CTRL versus KO ABX p=0.037, P45 WT CTRL versus KO CTRL p=0.372, P45 WT CTRL versus KO ABX p=0.907, P45 KO CTRL versus KO ABX p=0.063). **(B)** Fecal DNA content (ng) normalized on stool weight (mg). (One-way ANOVA p=0.008, multiple comparisons Tukey post-hoc test; WT CTRL versus KO CTRL p=0.624,

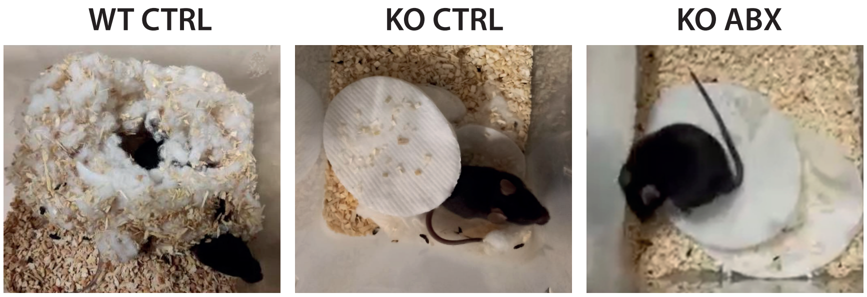


**SUPPL. Fig. 5.** Representative images of nest buildings. Pictures were taken 24h following nestlet placing.
